# Detection of genomic alterations in breast cancer with circulating tumour DNA sequencing

**DOI:** 10.1101/733691

**Authors:** Dimitrios Kleftogiannis, Danliang Ho, Jun Xian Liew, Polly Poon, Anna Gan, Raymond Chee-Hui Ng, Benita Kiat-Tee Tan, Kiang Hiong Tay, Swee-Ho Lim, Gek San Tan, Chih Chuan Shih, Tony Lim, Ann Siew-Gek Lee, Iain Tan, Yoon-Sim Yap, Sarah Ng

## Abstract

Analysis of circulating cell-free DNA (cfDNA) data has opened new opportunities for characterizing tumour mutational landscapes with many applications in genomic-driven oncology. We developed a customized targeted cfDNA sequencing approach for breast cancer (BC) using unique molecular identifiers (UMIs) for error correction. Our assay, spanning a 284.5 kb target region, is combined with freely-available bioinformatics pipelines that provide ultra-sensitive detection of single nucleotide variants (SNVs), and reliable identification of copy number variations (CNVs) directly from plasma DNA. In a cohort of 35 BC patients, our approach detected actionable driver and clonal SNVs at low (~0.5%) frequency levels in cfDNA that were concordant (83.3%) with sequencing of primary and/or metastatic solid tumour sites. We also detected *ERRB2* gene CNVs used for HER2 subtype classification with 80% precision compared to immunohistochemistry. Further, we evaluated fragmentation profiles of cfDNA in BC and observed distinct differences compared to data from healthy individuals. Our results show that the developed assay addresses the majority of tumour associated aberrations directly from plasma DNA, and thus may be used to elucidate genomic alterations in liquid biopsy studies.

## Introduction

One of the key objectives in precision oncology is to deliver better cancer diagnosis and tailored treatment. So far, analysis of tissue biopsy data is widely used to characterize tumour genomic landscapes, and to identify actionable somatic alterations^1,2^. However, tissues biopsies are invasive with constraints on frequency of tissue sampling, and may not be representative of the entire tumour load^3^.

As an alternative, recent studies have demonstrated the translational potentials of circulating cell-free DNA (cfDNA), or circulating tumor DNA (ctDNA) in cancer patients, for improving cancer management^3,4^. Such liquid biopsy data measured directly from body fluids (e.g. plasma) can be used to detect tumour somatic alterations, with the ability to provide early prognostication and/or better molecular profiling of patients with cancer without the risk and discomfort of invasive biopsies^5^. For example, estrogen receptor 1 (*ESR1*) mutation detected in the ctDNA of breast cancer (BC) patients pre-treated with aromatase inhibitors correlates with inferior treatment outcome on exemestane, but not on fulvestrant ^6,7^. *PIK3CA* mutation status based on ctDNA has also been demonstrated to predict benefit from *PI3K* inhibitor therapy in BC^8,9^. In other cancer types such as lung, ctDNA testing for *EGFR* mutation status has been approved by the Food and Drug Administration (FDA) to guide selection of therapy.

Several genetic techniques including digital droplet PCR (ddPCR) and BEAMing have been extensively applied to detect single nucleotide variations (SNVs) in cfDNA with very high precision (e.g. detection of alleles at lower than 0.1% frequency), but the analysis is restricted only to a limited number of genomic loci^5^. More recently, improvements in next generation sequencing (NGS) have allowed screening of broader genomic regions and simultaneous monitoring of multiple tumour-specific alterations in a single assay^2,10–12^. However, analyses of tumour NGS data from cfDNA is challenging due to several biological reasons (e.g. low cfDNA abundance in the blood stream) and other technical artifacts (e.g. error rates of NGS) that restrict the analytical sensitivity of tumour detection in plasma DNA.

In this study, we evaluate our targeted cfDNA sequencing approach that uses molecular barcodes for error correction. The developed assay spanning 77 genes (285.4 kb target region) is customized for BC, with focus on the commonly altered genes in BC as well as those with potential actionability^13,14^. To improve variant calling the developed assay is combined with freely available bioinformatics pipelines (https://github.com/dkleftogi/cfDNA_AnalysisPipeline) that enable ultra-sensitive detection of SNVs, accurate identification of copy number variations (CNVs) and evaluation of fragmentation profiles in cfDNA. We applied the developed cfDNA assay to detect genomic alterations in a cohort of 35 BC patients, and assessed the concordance of mutation calls with matched solid tumour sequencing.

## Materials and Methods

### Patient recruitment and sample collection

Patients were recruited at National Cancer Centre Singapore in a prospective observational study approved by the Singhealth Centralised Institutional Review Board (2013/251/B and 2014/119/B) where blood specimens were collected from patients with breast cancer from 2014 to 2016. Signed informed consent was obtained from all patients. From this study, we selected 30 metastatic cases with paired primary and\or metastatic specimens in addition to plasma taken prior to commencement of a new line of palliative systemic therapy (all subtypes), plus 5 patients (3 stage III, 2 stage II) about to commence neoadjuvant systemic therapy. For all patients recruited in the study buffy coat was also available for sequencing.

Retrospective review of medical and pathology records was performed to collect clinicopathologic details including patient demographics, tumour subtype via clinical testing, disease burden, and serum CA-15.3 level where available. The determination of estrogen receptor (ER), progesterone receptor (PR) and human epidermal growth factor receptor 2 (HER2) status by immunohistochemistry in this study was based on the latest recommendations by the American Society of Clinical Oncology and the College of American Pathologists^15,16^. ER and/or PR positive tumours that were HER2 negative were classified as hormone receptor positive (HR+)HER2-. Tumours with null expression in ER/PR and HER2 were classified as triple-negative breast cancer (TNBC) subtype. Tumours with positive HER2 expression (regardless of ER/PR status) were classified as the HER2-positive subtype. The demographic characteristics and clinical information for all patients included in the study are presented in Table 1.

**Table 1:** Clinicopathologic characteristics of study cohort.

| **Characteristic** | **Number**<br>**(n=35)** |
| --- | --- |
| Median age at study entry, in years | 50 (36-75) |
| <u>Ethnicity</u> |  |
| Chinese | 23 (65.7%) |
| Malay | 9 (25.7%) |
| Indian | 1 (2.9%) |
| Others | 2 (5.7%) |
| <u>Subtype</u> |  |
| HR+/HER2- | 18 (51.4%) |
| HER2+ (regardless of HR) | 14 (40.0%) |
| TNBC | 3 (8.6%) |
| <u>Stage</u> |  |
| II | 2 (5.7%) |
| III | 3 (8.6%) |
| IV (all relapsed after prior non-metastatic diagnosis) | 30 (85.7%) |
| Number of prior systemic therapies (including non-metastatic) |  |
| 0 | 5 (14.3%) |
| 1 or more | 30 (85.7%) |
| <u>Volume of Disease</u> |  |
| Low | 10 (28.6%) |
| Moderate | 14 (40.0%) |
| High | 11 (31.4%) |
| <u>Serum CA-15.3 levels</u> |  |
| Not raised | 15 (42.9%) |
| Raised | 18 (51.4%) |
| Not available | 2 (5.7%) |

Plasma samples from healthy individuals were collected similarly, as part of a parallel study (2012/733/B), where individuals were considered healthy if they were not cancer patients at time of collection. Aliquots of 1-2 ml of plasma were used for this study.

### Sample preparation and sequencing

Patient plasma was separated from whole blood within 6 hours of collection and subsequently frozen at −80°C. Plasma DNA was extracted using the QiaAmp Circulating Nucleic Acids kit (Qiagen). DNA was extracted from FFPE, frozen tissue and buffy coat samples using standard protocols. All DNA libraries were prepared using the Kapa Hyper Prep Kit (Kapa Biosystems, now Roche) using library adapters with a random 8-mer proximal to the library index site (IDT). Hybridization capture was done using an IDT Xgen Custom Panel of 77 genes (Supplementary Table 1) and reagents as per manufacturer’s instructions. Sequencing was performed on an Illumina Hiseq4000 (2 x 150 paired). Panel genes were chosen based on literature review for relevance and/or potential actionability in breast and other cancers.

### Processing of targeted sequencing cfDNA data

Raw sequencing data with unique molecular identifiers (UMIs) were demultiplexed using in-house pipelines. FASTQ files from different lanes were aligned to hg38 reference genome using bwa-mem^17^ (version 0.7.15), and merged to a single BAM file using GATK (version 4.1.1). UMI deduplication was performed using fgbio package (version 0.8.1). Reads with the same UMI were grouped together allowing one base mismatch between UMIs with minimum mapping quality 20. Consensus sequences were generated using the “adjacency” function by discarding groups of reads with single members^18^. Quality statistics and on-target analysis of coverage was performed using samtools^19^ (version 1.3.1) and bedtools^20^ (version 2.18).

### Bioinformatics pipeline for ultra-sensitive detection of SNVs in cfDNA

To identify SNVs in cfDNA we developed a bioinformatics pipeline that combines variant screening using VarDict^21^ with *in-silico* identification of duplexes using duplexCaller^12^. duplexCaller defines duplexes as pairs of read families (i.e. with different UMIs) that are mapped to the same genomic coordinates (start and stop) but with complementary sequencing orientation (i.e. one family in the forward direction and the other family in reverse). Using high-depth sequencing data with UMIs, duplex configuration approximates double stranded DNA molecules and guarantees reduction of false positive SNVs without sacrificing specificity^12^. To assess the detection capabilities of the proposed strategy in our cohort, we first performed a pilot analysis using data from healthy individuals. Under the assumption that low VAF SNVs in healthy data are more likely to be false positives and/or reflect problematic genomic positions, we performed manual inspection of variants below 2% VAF and found that the selection of the following parameters:

a. minimum VAF=0.005,
b. minimum base quality=30,
c. minimum coverage=100,
d. minimum number of supporting reads=3,
e. minimum number of supporting reads in forward strand=1,
f. minimum number of supporting reads in reverse strand=1,
g. number of duplexes=1,
h. minimum signal to noise ratio=20, and
i. mean position of reads supporting the variant greater than 15,

eliminates the majority of low VAF predictions in healthy data without affecting the true positive rate (i.e. variants at VAF > 10% are unaffected). For small deletions/indels we applied the same criteria with more stringent values for minimum VAF=0.01, minimum coverage=200, minimum number of supporting reads=6, signal to noise ratio=25, number of duplexes=2. We also filtered out recurrent variants that fall into high discrepancy genomic regions (HDR) as described in Bernell et al.^22^ and blacklisted these sites in further analyses.

Based on the aforementioned criteria we performed variant calling in cfDNA samples from BC patients. To identify somatic variants in cfDNA samples, we required a) zero support of reads in the matched normal (buffy coat) sample, with minimum coverage of at least 50 reads in the normal, and b) that the variant be absent from databases of common polymorphisms eg. 1000 Genomes project phase 3. Finally, the list of somatic variants per patient was annotated using Ensembl VEP ^23^, and SNVs (including small indels/deletions) with MODERATE or HIGH impact were considered for further analyses.

### Processing and SNV calling of sequencing data from solid tumour sites

Targeted sequencing data from solid tumour sites were aligned to hg38 reference genome using bwa-mem (version 0.7.15). Data from different lanes were merged using GATK version 4.1.1 and duplicates were removed using GATK MarkDuplicates function. Quality statistics and on-target analysis of coverage was performed using samtools (version 1.3.1) and bedtools (version 2.18) (Supplementary Table 2).

To identify SNVs in solid tumour samples without UMIs, we applied a previously developed variant calling pipeline^24^ that combines the Mutect2 variant caller^25^ with Platypus^26^. Mutect2 was first run with default parameters on tumour and normal samples of every patient. Then, we used the VCF files returned by Mutect2 as priors to Platypus with zero posterior probability, and jointly called variants. To identify somatic variants in solid tumour samples, we required a) minimum coverage of 50 reads, b) at least 2 reads supporting the variant, c) zero supporting reads in the matched normal sample with minimum coverage of 50 reads, d) quality filtering flag returned by Platypus either ‘PASS’, ‘Q20’, ‘QD’, ‘alleleBias’ or ‘HapScore’, and e) absence from the list of common polymorphisms reported in the 1000 genomes database.

### Identification of Copy Number Variations (CNVs)

CNVs were called in ctDNA and solid tumour samples using CONTRA with the Null Distribution Estimation (NDE) workflow^27^ using a multimodal distribution instead of the unimodal distribution used by default. We took the best fit between a bimodal and trimodal distribution as determined by the Akaike information criterion (AIC) estimator and fed the model parameters into the software’s threshold cutoffs for CNV identification. All other parameters of CONTRA remained unchanged.

### Estimation of cfDNA fragmentation profiles

We developed a bioinformatics pipeline to characterize fragment length profiles of cfDNA samples. The pipeline takes as input BAM files deduplicated with UMIs and uses pysam libraries (https://pysam.readthedocs.io/en/latest/index.html) to extract the fragment length values based on the TLEN sam flag of all read pairs mapped to the target region. We considered read pairs with minimum mapping quality 10, and excluded read pairs where mates were mapped to different chromosomes. The observed data were binned and normalized by the total number of read pairs sequenced in the sample. Following this procedure, we generated density profiles for all BC patients and healthy individuals. All reads from all healthy individuals were pooled to generate a combined fragment length profile that was used as a reference.

### Code availability

The bioinformatics workflow described in this study can be downloaded from our GitHub repository (https://github.com/dkleftogi/cfDNA_AnalysisPipeline) with an overview given in Supplementary Figure 1. We provide a collection of scripts written in Python for UMI-aware BAM file deduplication, mutation calling and fragment length analysis as well as a Conda virtual environment to resolve dependencies with required packages.

## Results

### SNVs detected in cfDNA samples

We first determined the feasibility of the developed cfDNA sequencing approach to detect tumour-specific alterations from cfDNA of patients with BC. Our assay combined with a customized bioinformatics pipeline for variant calling provides sufficient sensitivity to detect tumour-associated SNVs in 30 out of 35 samples with median VAF of 0.02 (Figure 1, Supplementary Table 3). Across 30 samples with detectable SNVs, a total of 98 variants were detected, with ~71% of them already reported in existing databases (e.g. ClinVar and/or COSMIC). The increased number of existing variants detected using the deployed cfDNA assay highlights the capability of identifying biologically important mutations (i.e., cancer-related or associated to other human disorders) directly from cfDNA. Among the 5 patients with no detectable tumour variants in cfDNA, 4 of them had low tumour burden based on clinical and radiological information (Table 1, Figure 1).

**Figure 1.**
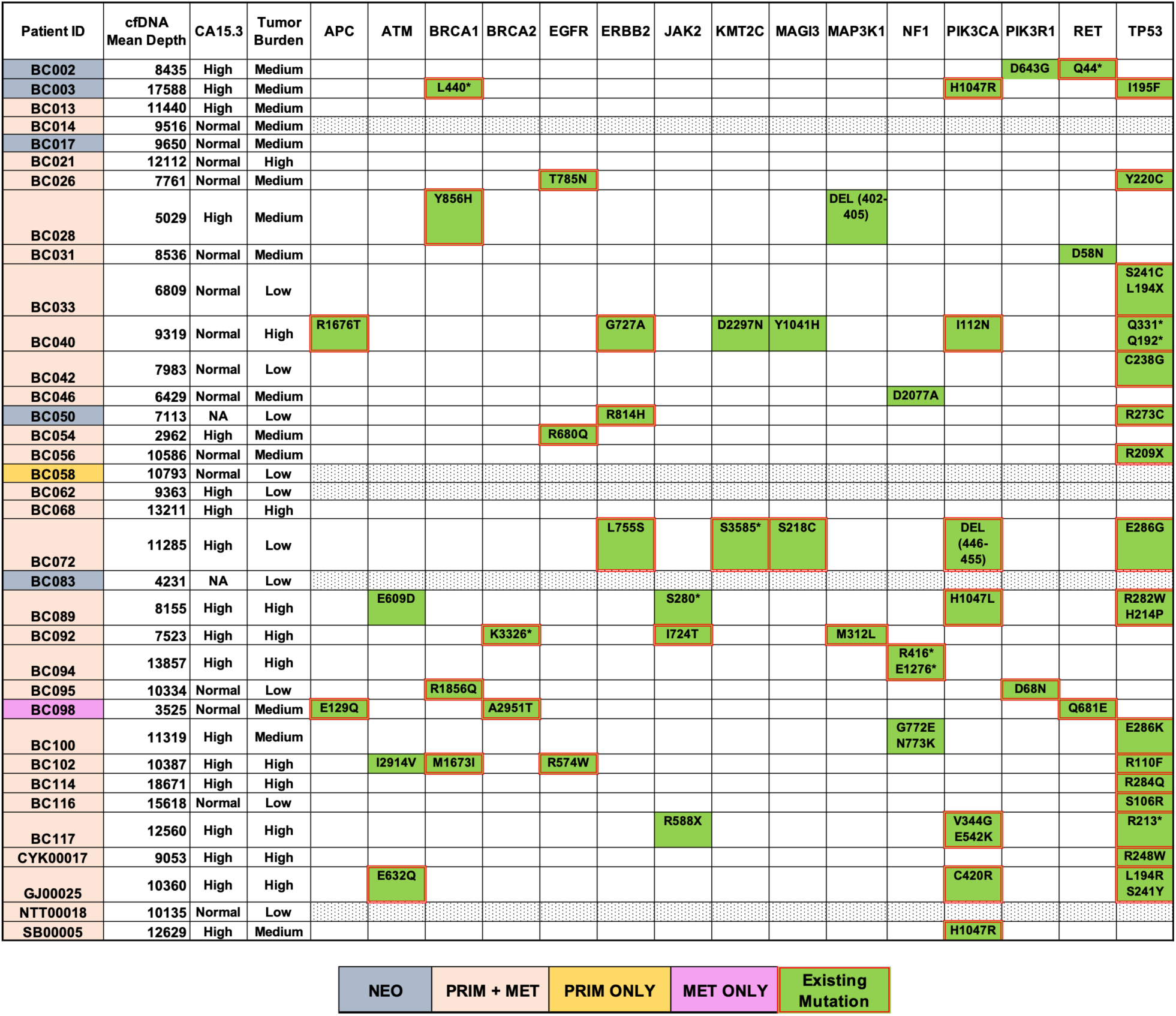
SNVs found in known cancer-related genes by cfDNA sequencing. 35 BC patients were sequenced using the developed cfDNA assay. One sample had only sequencing of cfDNA and the primary site available (orange), one with only cfDNA and the metastatic site (pink) and five were neoadjuvant samples (grey). All others had cfDNA, primary and metastatic sites sequenced. Variants that are already in known databases are outlined in red. Median cfDNA sequencing depth, CA15.3 level and assessed tumour burden is also listed.

Figure 1 summarizes known tumor suppressor or oncogenic genes such as *TP53* (found mutated in 16/35 patients), *PIK3CA* (7/35), *BRCA1* (4/35), *NF1* (3/35), *EGFR* (3/35), *ERBB2* (3/35), *ATM* (3/35) or *RET* (3/35) that were recurrently mutated in our cohort. Overall 18 out of 25 patients harbored more than two mutated genes, likely associated with parallel cancer evolution, and/or resistance mechanisms to combinatorial chemotherapy.

To investigate the translational potential of the identified variants in cfDNA we performed correlation analysis using the cancer antigen 15-3 (CA15-3) tumour marker, which is frequently used in routine clinical practice. We computed the median VAF of all patients in the cohort and stratified them based on their tumour burden based on clinical and imaging reports. We found that patients with high volume of disease (such as patients with widespread metastatic disease or in visceral crisis), harbor variants in cfDNA at much higher VAF levels compared to tumours of low (clinical stage or oligometastatic disease) and medium burden (burden of disease intermediate between high and low disease burden) (Figure 2a, Wilcoxon rank sum test p value = 4.56e-04). We also found that patients with high levels of the CA15-3 harbor mutations in cfDNA at higher VAF levels compared to tumours of normal CA15-3 levels (Figure 2b, Wilcoxon rank sum test pvalue=0.02).

**Figure 2.**
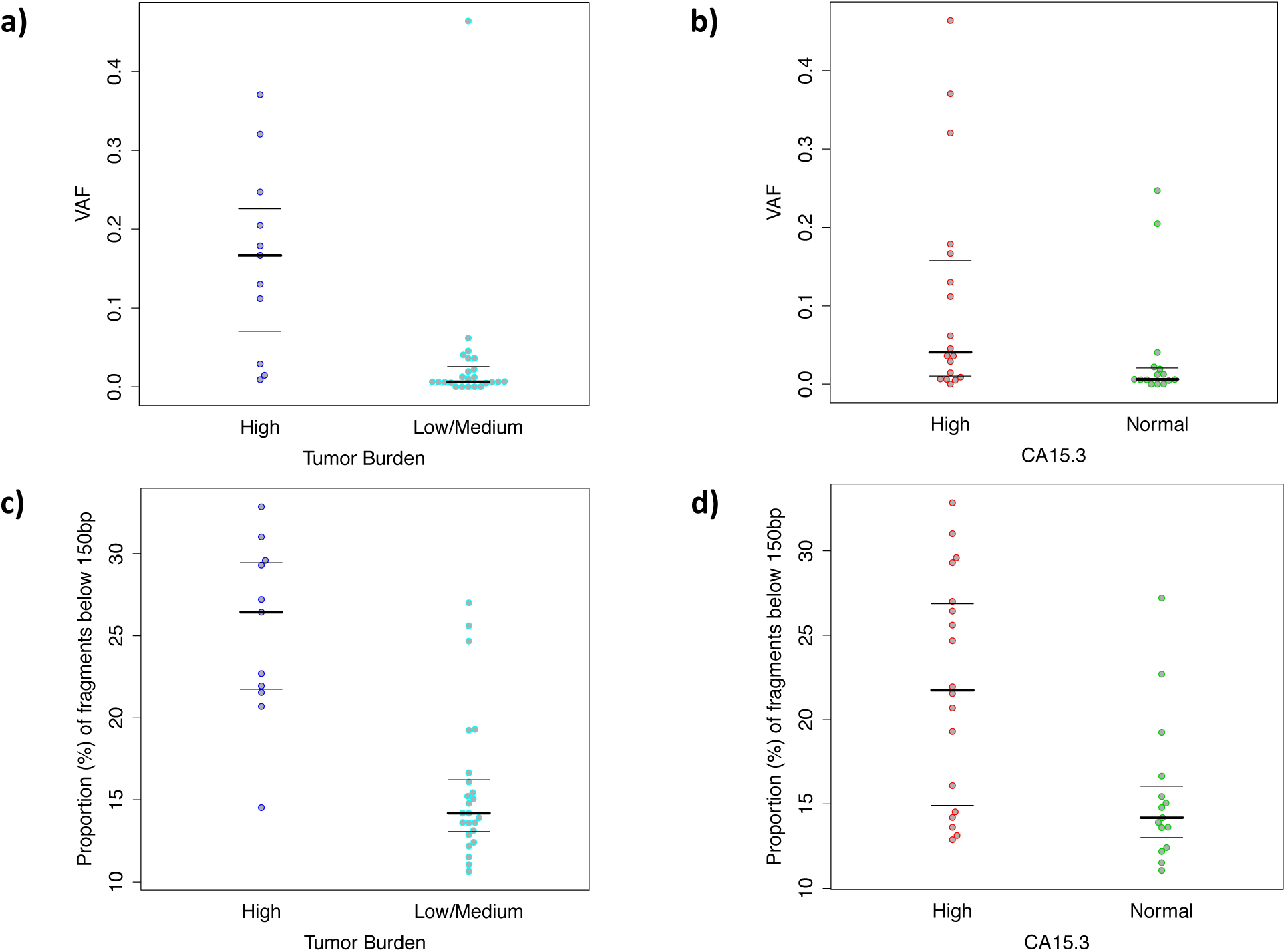
Association between genomic alterations detected in cfDNA and clinical information. Our analysis shows that the median VAF of SNVs detected in cfDNA as well as the proportion of shorter fragments of cfDNA correlates well with tumour burden and serum CA15.3 marker. Shown are the median VAF levels detected in cfDNA in patients with a) High and Low/Medium tumour burden, and High and Normal CA15.3 levels, as well as the proportion of cfDNA fragments below 150bp for patients with c) High and Low/Medium tumour burden and d) High and Normal CA15.3 levels.

### Concordance of cfDNA and sequencing of solid tumour sites

Out of the 30 metastatic BC cases, 28 patients had available sequencing from matched cfDNA, primary tumour and metastatic sites. We utilized this subset of samples to assess the concordance and discordance of variants detected in cfDNA. Our cfDNA assay detected 84 mutations across these plasma samples. Coverage statistics, number of supporting reads and VAF for all variants detected in cfDNA are shown in Supplementary Figure 2. Overall, we found discrepancies between the data quality of cfDNA, primary and metastatic tumour sites attributed to differences in DNA material and sequencing protocols applied in the study (i.e. ultra-deep sequencing for cfDNA with UMIs, and much lower depth sequencing without UMIs for DNA from FFPE samples).

We restricted our analysis in 54 out of 84 positions that were “callable” in both solid tumour sites (i.e. coverage of more than 50 reads in primary and metastatic samples). Using this set of 54 callable positions we identified 25 (~46.3%) mutations that were concordant between cfDNA, primary and metastatic samples, Figure 3 shows the VAF levels of all variants validated by sequencing in the cohort. We also detected 12 mutations (~15% of positions tested) with VAF in cfDNA as low as 0.01, but with higher VAF at the matched solid tumour sites. Such examples highlight the effectiveness of the developed assay to monitor important cancer alterations directly from cfDNA, even when mutation-bearing molecules are very rare in plasma.

**Figure 3.**
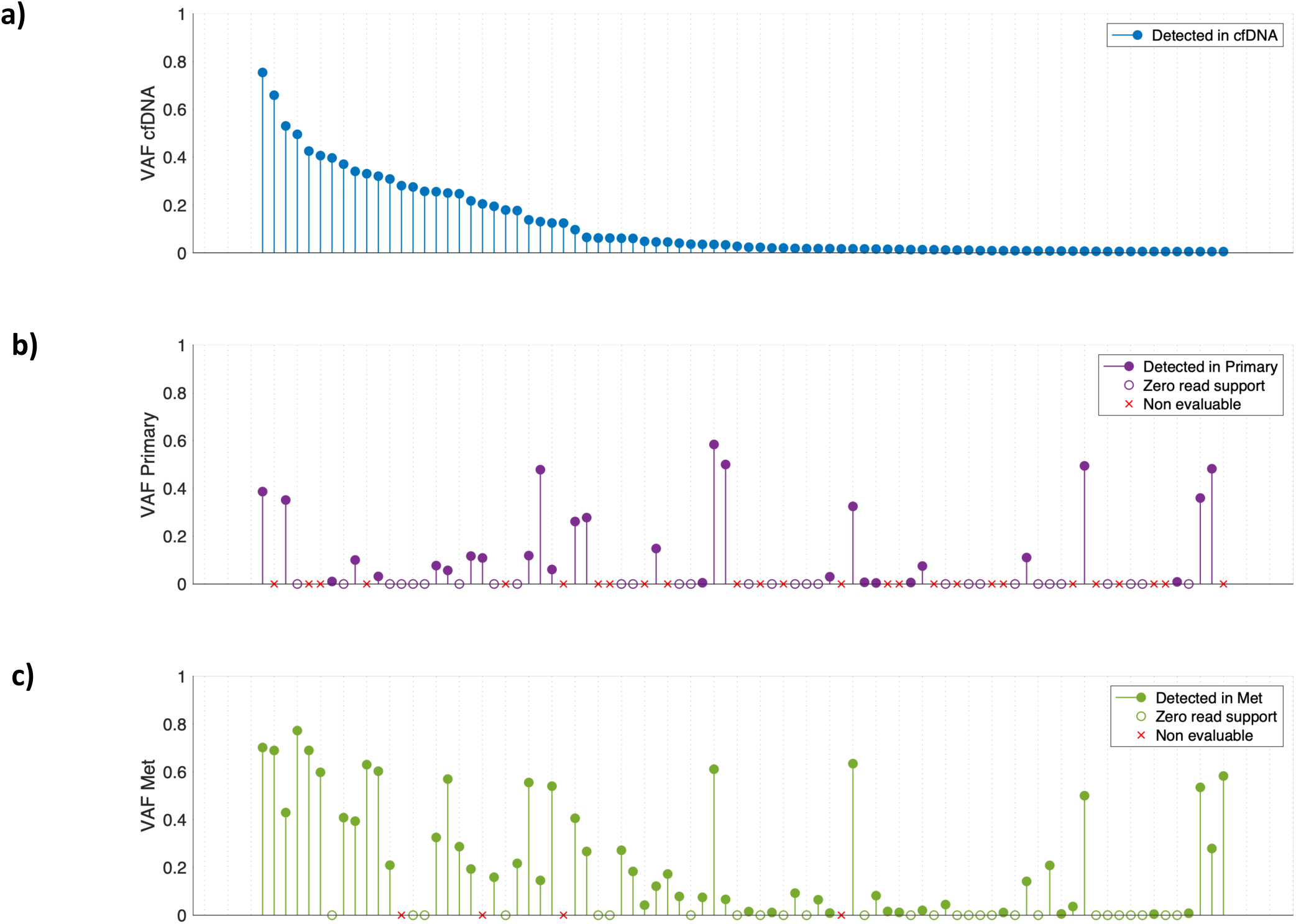
Concordance of SNVs identified by cfDNA sequencing and by sequencing of solid tumour sites. The a) cfDNA VAF, b) VAF in the primary site and c) VAF in the metastatic site are displayed for the 84 SNVs detected in cfDNA for patients with available sequencing for all of cfDNA, primary and metastatic tumour. In a), the minimum VAF was set at 0.005. In b) and c), positions that did not achieve 50X coverage are marked with ‘x’, while positions that did but nonetheless had 0 reads corresponding to the alternative allele are marked with empty circles.

Next, we relaxed the conditions and we analyzed the concordance of variants detected in cfDNA to at least one of primary or metastatic samples. Using this approach we identified 45 (~83.3%) variants that were concordant between cfDNA and one of the two solid tumour sites. Figure 3 shows that the majority of concordant variants have higher VAF in metastatic sites, which is consistent with the patient clinical progression at time of sampling. However, the results of this analysis should be interpreted with caution, because while emergence of new variants in metastatic tumours is not unexpected, we cannot completely rule out that the absence of variants in primary sites is due to technical reasons, including inferior sample quality of FFPE-extracted DNA from solid tumour sites.

Then, we assessed SNVs called in solid tumour sites and not in cfDNA, for which there were only seven variants. From them, two SNVs (genes *PIK3CA* and *TP53*) detected in patients with non-metastatic disease, two SNVs (genes *MAGI3* and *EZH2*) were detected in the primary tumour samples of two patients, and three SNVs (genes *APC, TP53* and *MDM4*) were detected in metastatic samples of other patients. Five out of seven SNVs not detected in cfDNA had zero support of the alternative alleles in cfDNA, except for two mutations (gene *TP53* position Y236R with 13 reads, and gene *MDM4* position A42V with 29 reads) that had sufficient supporting reads, but they were not supported by duplexes. Such cases might represent genuine false negatives of our variant calling pipeline.

Together, our data show that cfDNA screening using the developed assay addresses the majority of SNVs found by solid tumour sequencing, including in BC patients with low tumor burden.

### *ERBB2* CNV analysis and HER2 subtype classification in BC

In this subsection we investigate whether we could use the developed cfDNA assay to detect CNVs in BC. We applied a customized version of the CONTRA algorithm to infer CNV profiles using the reads generated by targeted hybrid capture cfDNA sequencing. We focused on the *ERBB2* (HER2) oncogene to identify detectable amplifications that may be used for HER2 subtype classification. For validation purposes, all patients in the cohort had previously undergone HER2 testing mainly via immunohistochemistry, with fluorescent in-situ hybridization (FISH) testing performed for equivocal immunohistochemical results. Fourteen of them were found to be HER2+, whereas 18 patients were HR+HER2- and 3 were Triple-Negative (TNBC). Our cfDNA approach detected correctly 12 of HER2+ patients (*ERBB2* amplified) and 18 of HER2- / TNBC patients, whereas it misclassified 5 patients. From the misclassified cases, 3 patients were falsely predicted to be HER2+, and 2 patients were falsely predicted to be HER2-, classification performance that could be translated to 80% positive predictive value (Precision), 85.7% true positive rate (Sensitivity) and 85.7% true negative rate (Specificity). Using the same algorithm we also generated *ERBB2* CNV profiles using sequencing data from primary and metastatic solid tumour sites (Figure 4). We note that the *ERBB2* amplification was also detected correctly in 10 out of 14 patients using data from matched metastatic tumour sites, and in 7 out of 14 patients using data from primary tumour sites. In total, our findings using orthogonal sequencing and immunohistochemistry validate further the ability of our cfDNA assay to identify *ERBB2* CNVs in BC without prior knowledge of tumour sequencing.

**Figure 4.**
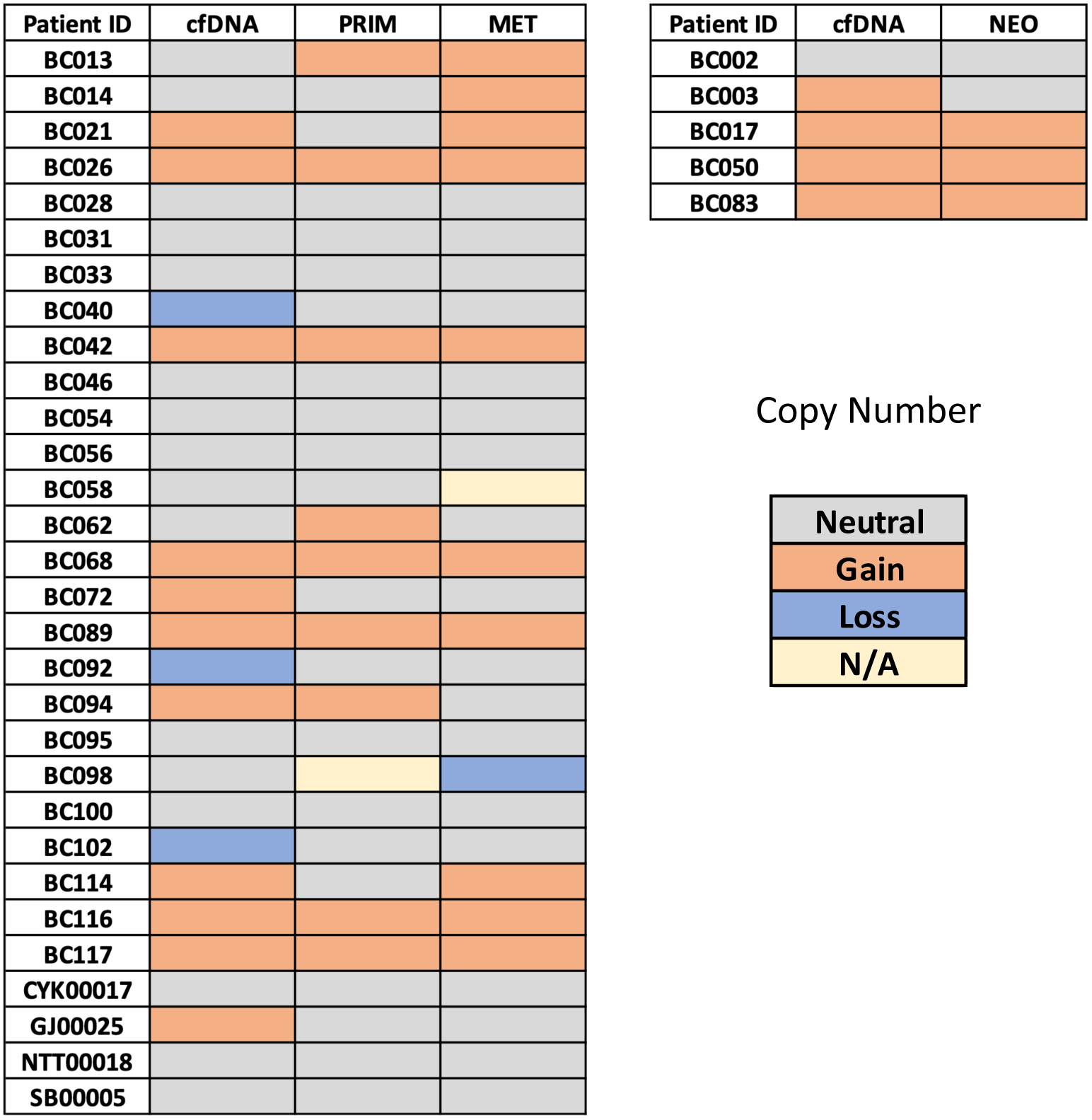
Identification of *ERBB2* gene CNVs. CNVs detected in *ERBB2* (HER2) gene using sequencing data from cfDNA, primary and metastatic solid tumour sites..

### cfDNA fragmentation profiles of BC patients

Recent analyses of the fragment sizes of cfDNA of patients with cancer, suggest altered fragmentation profiles compared to healthy individuals ^28^. Here, we used our developed cfDNA assay to characterize the fragment length profiles of 35 patients with BC and to explore its potential as a biomarker for disease monitoring. For comparison purposes, we also analyzed 38 samples from 19 healthy individuals (two technical replicates with different library preparations) and compiled a reference fragment length profile. Figure 5a shows the combined fragmentation profile of healthy individuals, and Figure 5b shows different profiles of patients with BC. Our analysis detected many differences in cfDNA profiles of patients with BC, whereas healthy profiles were much more similar to each other (Figure 5c Wilcoxon rank sum test p-value= 6.7608e-12). Our data confirmed that the median fragment length of cfDNA of patients with cancer is also different from healthy individuals (Figure 5d, Wilcoxon ranksum test pvalue= 2.8000e-05). However, the differences between the median fragment size of healthy individuals (overall median 167 bp) and fragment size of patients with cancer (overall median 166 bp) were only few bases in size. More interestingly, we estimated that the proportion of fragments below 150 bp is much higher in BC (Figure 5e, Wilcoxon ranksum test p-value= 1.4060e-12) compared to the healthy reference. Our observations about the proportion of fragments below 150 bp are in concordance with the results from other studies, using mainly low-pass WGS samples from treatment naïve or early stage patients^28,29^.

**Figure 5.**
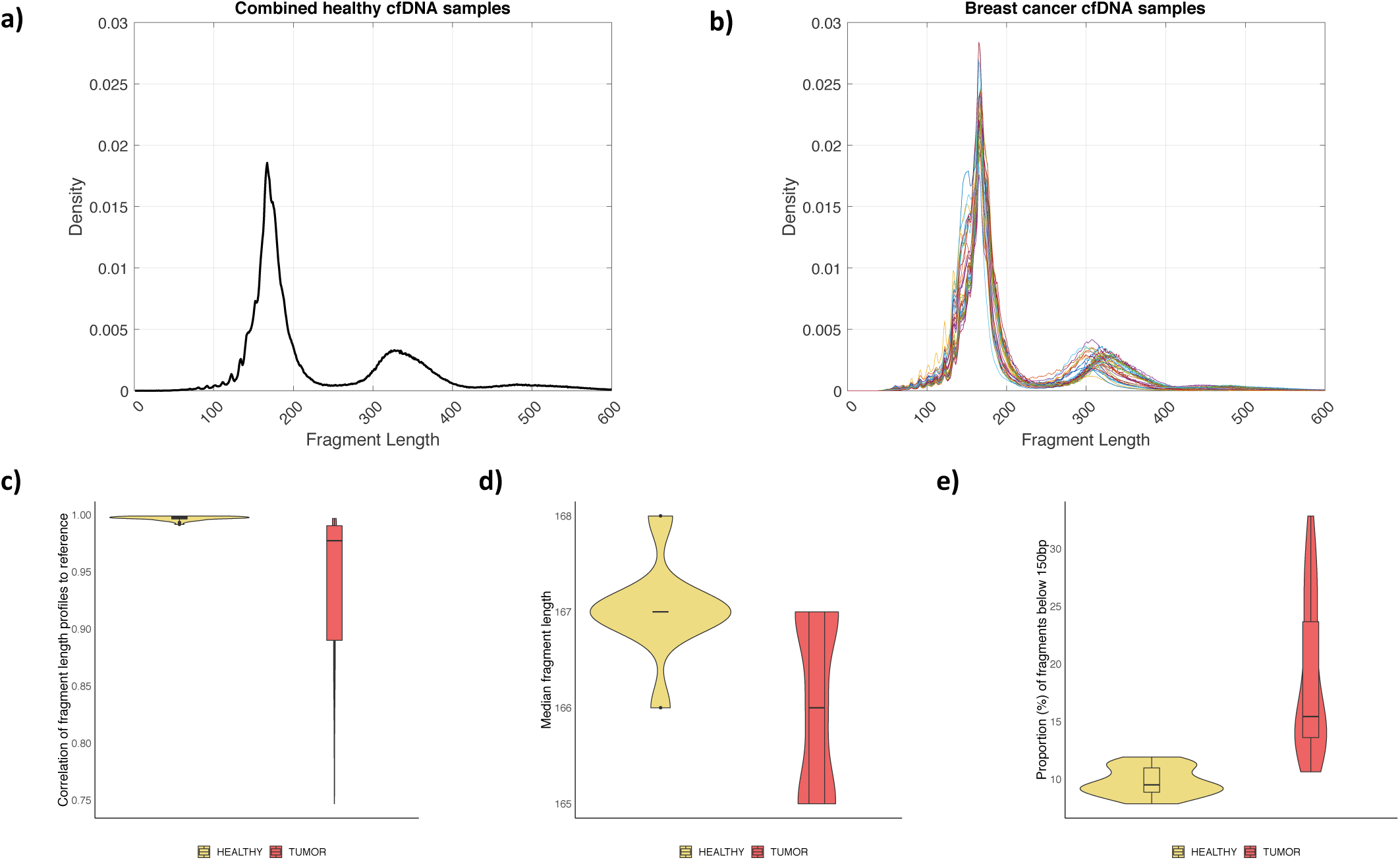
Analysis of cfDNA fragmentation profiles in BC. a) Reference cfDNA fragmentation profile complied using data (two technical replicates) from 19 healthy individuals. b) cfDNA fragmentation profiles of 35 patients with BC included in the study. c) cfDNA fragmentation profiles of 19 healthy individuals are highly correlated to the reference profile, whereas cfDNA fragmentation profiles of 35 BC patients have much lower correlation to the reference healthy profile. d) Median cfDNA fragment sizes in bp are shown for 19 healthy individuals and 35 BC patients. e) The proportion of short cfDNA fragments (below 150bp) detected in 19 healthy individuals is much lower compared to the proportion of short fragments detected in 35 BC patients.

These results indicate that fragmentation profiles of cfDNA in BC integrate genomic and epigenomic (i.e. nucleosome positioning) features that could serve as novel biomarkers in clinical settings. To investigate this hypothesis in our cohort, we associated our findings with CA15-3 levels and tumour burden, which was assessed as above. We found that patients with high tumour burden have a much higher proportion of shorter fragments compared to patients with low or medium tumour burden (Figure 2c, Wilcoxon ranksum test pvalue=1.33e-04). We also found that shorter fragments are also proportionally higher in patients with elevated CA15.3 levels, compared to patients with normal CA15-3 levels (Figure 2d, Wilcoxon ranksum test pvalue= 0.0088). Consequently, our findings based on fragmentation profiles are in concordance with the results obtained from routine cancer biomarkers.

Then, we explored the potential of the fragment length analysis to monitor patients’ disease status. We used the median VAF levels in cfDNA as a proxy to disease burden, and we measured the deviation of the corresponding fragment length profiles to the healthy reference. We observed (Figure 6a) that the deviation of fragment length profiles of patients with BC, follows the VAF levels with high correlation (Pearson’s correlation coefficient 0.792). This observation opens possibilities for the development of new biomarkers for better disease management that could complement variant analysis based on VAFs.

**Figure 6.**
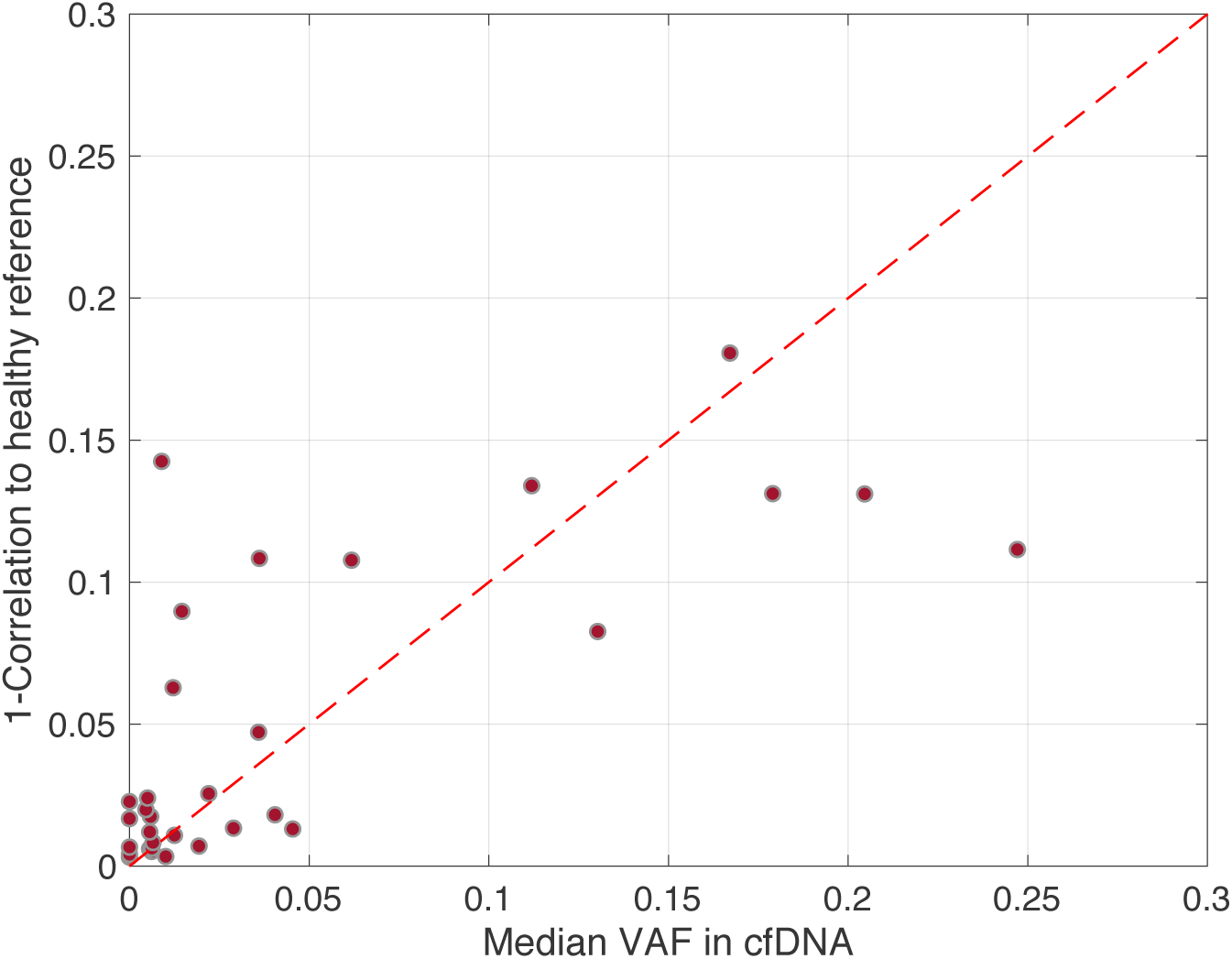
Association between cfDNA fragmentation profiles and median VAF levels detected in cfDNA. We show that the variation of cfDNA fragmentation profiles of patients with BC compared to the healthy reference (measured as 1 - Pearsons correlation between the two profiles), correlates well (Pearson’s correlation 0.793) with the median VAF levels of SNVs detected in cfDNA.

## Discussion

Several recent studies have demonstrated the ability of cfDNA sequencing to provide early prognostication, better molecular profiling and monitoring of disease dynamics. We developed a new error-corrected cfDNA sequencing approach customized for BC, in combination with publicly available bioinformatics strategies for the identification of tumour-associated genomic alterations. The developed NGS assay covers 77 known cancer-related genes (285.4 kb target region), providing the opportunity to elucidate genomic alterations with significant clinical value without prior knowledge of tissue sequencing.

Usage of UMIs for error correction, combined with SNV calling based on *in-silico* identified duplex DNA molecules, allowed us to detect low VAF SNVs in ctDNA samples with reduced number of false positives. In a proof-of-concept analysis using clinical data from 35 BC patients, the developed assay detected SNVs in 30 out of 35 cfDNA samples with ~71% of variants already reported in known databases. The majority of detected mutations in cfDNA were concordant with sequencing of solid tumour sites. However, cfDNA sequencing identified more potential mutations, many with VAF below 1%, highlighting the ability of our assay to track rare mutations in plasma. This opens possibilities for longitudinal monitoring of cancer-genomic alterations directly from plasma, without the increased risk and cost of invasive needle biopsies.

The presence of SNVs called only in ctDNA, although not-validated using orthogonal approaches (e.g. ddPCR), is consistent with tumour evolution hypotheses, where multiple low VAF clonal and sub-clonal alterations may be the basis of resistance to several lines of chemotherapy. These sites are usually not detectable in solid tumour sequencing due to lack of read depth. Importantly, our approach for SNV detection does not rely on statistical modelling for background noise estimation^12,30^. Such approaches require big cohorts of reference data (e.g. healthy) that are usually difficult to collect, and may not be easily extended or generalized (e.g. if the panel size increases). In contrary our approach relies on a number of adjustable post-filtering criteria, it is much simpler, faster and thus it can serve as a paradigm for subsequent studies.

Our assay was also effective on the detection of CNVs in important oncogenes such as *ERBB2* that are equally important for understanding tumorigenesis and deciphering tumor progression mechanisms. This opens opportunities for better tumor characterization, where sequential plasma samples can be collected to portray more accurately CNVs and SNVs across time.

Finally, our study reports on cfDNA fragmentation profiles in BC. Our data from 35 BC patients and 19 healthy individuals re-confirmed that cfDNA fragmentation profiles recapitulate genomic and epigenomic features that are in principle, different. Interestingly, we found that in patients with BC the deviation of cfDNA fragmentation profile from the healthy reference correlates well with the mutational load measured by VAF in cfDNA. This observation opens new avenues for monitoring patients’ progression under treatment, and developing cancer biomarkers using cfDNA sequencing. Towards this, future investigations are required to establish whether the determination of cfDNA fragmentation profiles might provide prognostic value for patients with early stage BC.

However, similar to all other cfDNA sequencing studies, our approach has several limitations. First, ultra-sensitive detection of cancer especially at early stage patients is not always viable^31^. Low abundance of cfDNA in plasma combined with increased errors of NGS might limit the applicability of the developed assay. We note that although our assay uses barcoded libraries, multiple sources of errors during library preparation and PCR amplification restrict the analytical sensitivity for SNV detection. For this reason, in our analysis with BC data, we used a 0.5% minimum VAF cutoff, after experimentation and manual inspection of healthy data without assessing our assay’s detection limits using analytical approaches. This part of our pipeline, might be improved in future studies when more control data (e.g. spike-in) become available. Nevertheless it is also possible that a few true variants are not supported by duplexes. We observed two mutations found in solid tumour sites that were filtered out in cfDNA samples due to absence of duplex support, indicating that there is room for further fine-tuning of parameters, using a larger training cohort.

In addition, the developed targeted assay is less powerful on the detection of structural variants (e.g. translocations or inversion) compared to whole exome or whole genome sequencing data. This is an inherent limitation of selective sequencing and further work may be done in the development of more accurate methods for CNV detection in targeted NGS data. Finally, the small cohort size and the absence of serial specimens for comparisons, limit our ability to associate the findings of this study with clinical outcome.

In conclusion, our data show that the developed assay combined with freely available bioinformatics approaches addresses the majority of tumour associated aberrations that could complement routine biomarkers (e.g., CA15.3 antigen) and other diagnostic tools in a minimally invasive liquid biopsy setup.

## Acknowledgements

The authors thank the patients and the healthy volunteers for their kind participation in the study, and the SingHealth Tissue Repository for providing selected fresh frozen tumor specimens.

## Supplementary Material

**Supplementary Table 1:** Seventy seven genes incorporated in the developed cfDNA assay

|  |  |  |  |  |
| --- | --- | --- | --- | --- |
| CDH1 | CDKN1B | EZH2 | KRAS | ERBB4 |
| CDKN2A | DOT1L | FLT3 | NRAS | FGFR1 |
| HRAS | ERBB3 | GNA11 | PIK3CA | FGFR2 |
| JAK2 | ESR1 | GNAQ | PIK3R1 | MET |
| JAK3 | GATA3 | GNAS | SMAD4 | FGFR3 |
| NF1 | JAK1 | HNF1A | TP53 | ALK |
| NOTCH1 | MAGI3 | IDH1 | FBXW7 | MAP2K1 |
| RB1 | MDM2 | IDH2 | RNF43 | MTOR |
| RET | MDM4 | KDR | APC | MAP2K4 |
| SRC | KMT2C | MPL | EGFR | MAP3K1 |
| STK11 | NOTCH4 | NPM1 | ZNRF3 | AKT1 |
| ARID1A | TSC1 | PTPN11 | ATM | PDGFRA |
| ATR | TSC2 | SMARCB1 | PTEN | KIT |
| AURKA | ABL1 | SMO | CTNNB1 |  |
| BRCA1 | CSF1R | VHL | MLH1 |  |
| BRCA2 | DDR2 | BRAF | ERBB2 |  |

**Supplementary Table 2:**
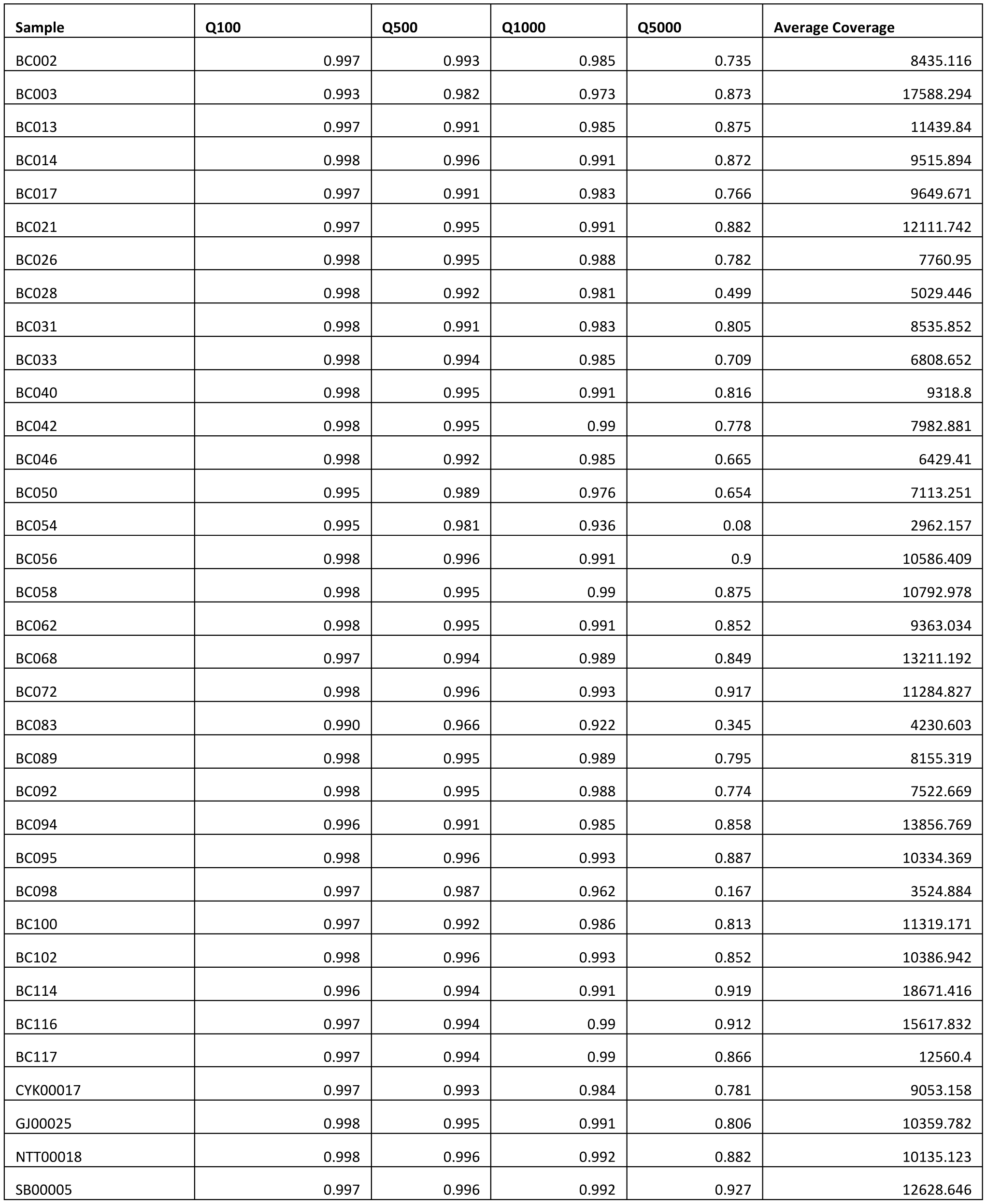
Analysis of coverage for all cfDNA samples included in the study cohort. In columns Q100, Q500, Q1000 and Q5000, we show the (%) fraction of positions in the panel covered by at least 100,500,1000 and 5000 reads. We also show the average coverage levels achieved across the panel.

| Sample | Q100 | Q500 | Q1000 | Q5000 | Average Coverage |
| --- | --- | --- | --- | --- | --- |
| BC002 | 0.997 | 0.993 | 0.985 | 0.735 | 8435.116 |
| BC003 | 0.993 | 0.982 | 0.973 | 0.873 | 17588.294 |
| BC013 | 0.997 | 0.991 | 0.985 | 0.875 | 11439.84 |
| BC014 | 0.998 | 0.996 | 0.991 | 0.872 | 9515.894 |
| BC017 | 0.997 | 0.991 | 0.983 | 0.766 | 9649.671 |
| BC021 | 0.997 | 0.995 | 0.991 | 0.882 | 12111.742 |
| BC026 | 0.998 | 0.995 | 0.988 | 0.782 | 7760.95 |
| BC028 | 0.998 | 0.992 | 0.981 | 0.499 | 5029.446 |
| BC031 | 0.998 | 0.991 | 0.983 | 0.805 | 8535.852 |
| BC033 | 0.998 | 0.994 | 0.985 | 0.709 | 6808.652 |
| BC040 | 0.998 | 0.995 | 0.991 | 0.816 | 9318.8 |
| BC042 | 0.998 | 0.995 | 0.99 | 0.778 | 7982.881 |
| BC046 | 0.998 | 0.992 | 0.985 | 0.665 | 6429.41 |
| BC050 | 0.995 | 0.989 | 0.976 | 0.654 | 7113.251 |
| BC054 | 0.995 | 0.981 | 0.936 | 0.08 | 2962.157 |
| BC056 | 0.998 | 0.996 | 0.991 | 0.9 | 10586.409 |
| BC058 | 0.998 | 0.995 | 0.99 | 0.875 | 10792.978 |
| BC062 | 0.998 | 0.995 | 0.991 | 0.852 | 9363.034 |
| BC068 | 0.997 | 0.994 | 0.989 | 0.849 | 13211.192 |
| BC072 | 0.998 | 0.996 | 0.993 | 0.917 | 11284.827 |
| BC083 | 0.990 | 0.966 | 0.922 | 0.345 | 4230.603 |
| BC089 | 0.998 | 0.995 | 0.989 | 0.795 | 8155.319 |
| BC092 | 0.998 | 0.995 | 0.988 | 0.774 | 7522.669 |
| BC094 | 0.996 | 0.991 | 0.985 | 0.858 | 13856.769 |
| BC095 | 0.998 | 0.996 | 0.993 | 0.887 | 10334.369 |
| BC098 | 0.997 | 0.987 | 0.962 | 0.167 | 3524.884 |
| BC100 | 0.997 | 0.992 | 0.986 | 0.813 | 11319.171 |
| BC102 | 0.998 | 0.996 | 0.993 | 0.852 | 10386.942 |
| BC114 | 0.996 | 0.994 | 0.991 | 0.919 | 18671.416 |
| BC116 | 0.997 | 0.994 | 0.99 | 0.912 | 15617.832 |
| BC117 | 0.997 | 0.994 | 0.99 | 0.866 | 12560.4 |
| CYK00017 | 0.997 | 0.993 | 0.984 | 0.781 | 9053.158 |
| GJ00025 | 0.998 | 0.995 | 0.991 | 0.806 | 10359.782 |
| NTT00018 | 0.998 | 0.996 | 0.992 | 0.882 | 10135.123 |
| SB00005 | 0.997 | 0.996 | 0.992 | 0.927 | 12628.646 |

**Supplementary Table 3.**
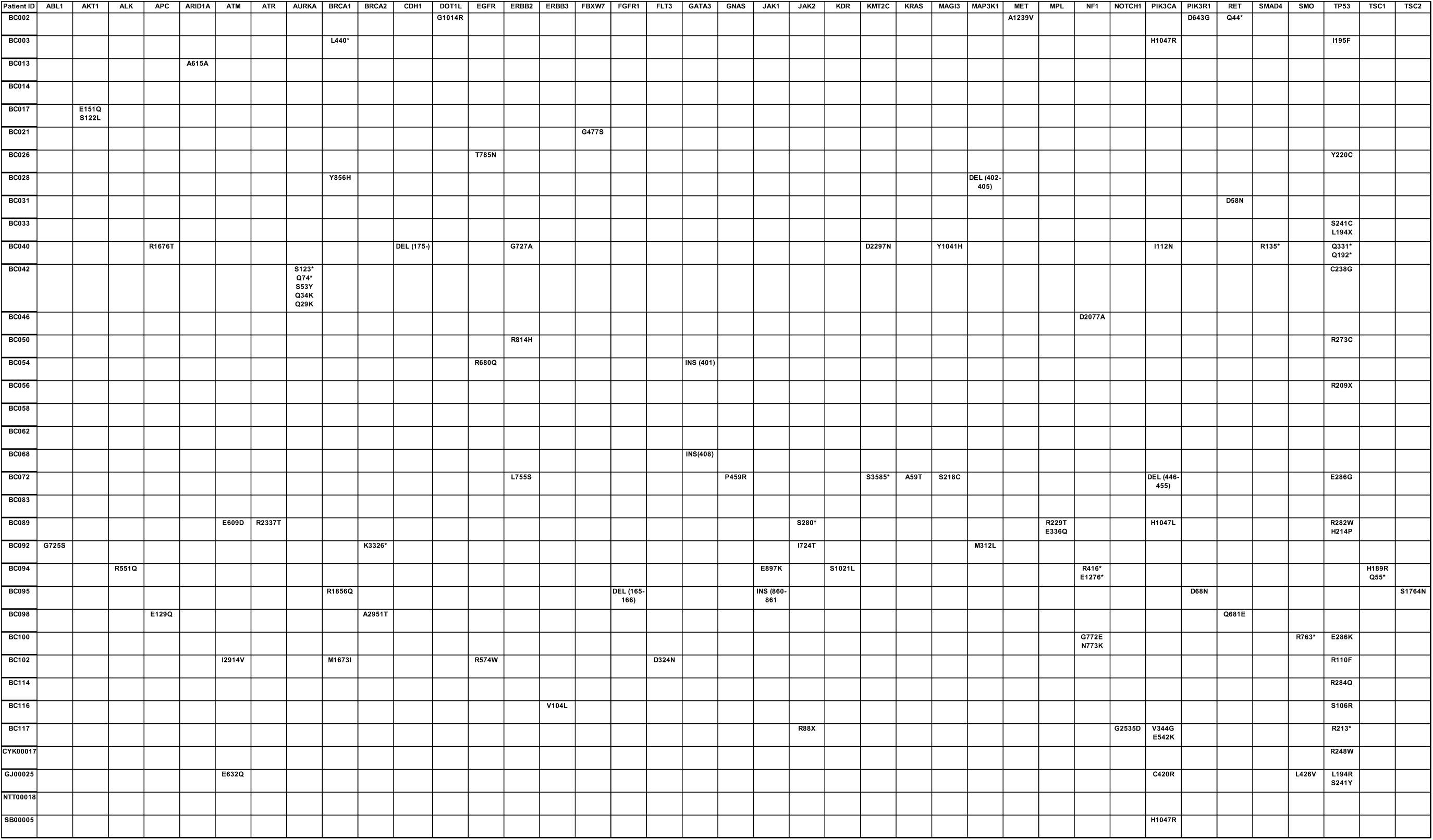
All SNVs identified by cfDNA sequencing in the study cohort.

**Supplementary Figure 1.**
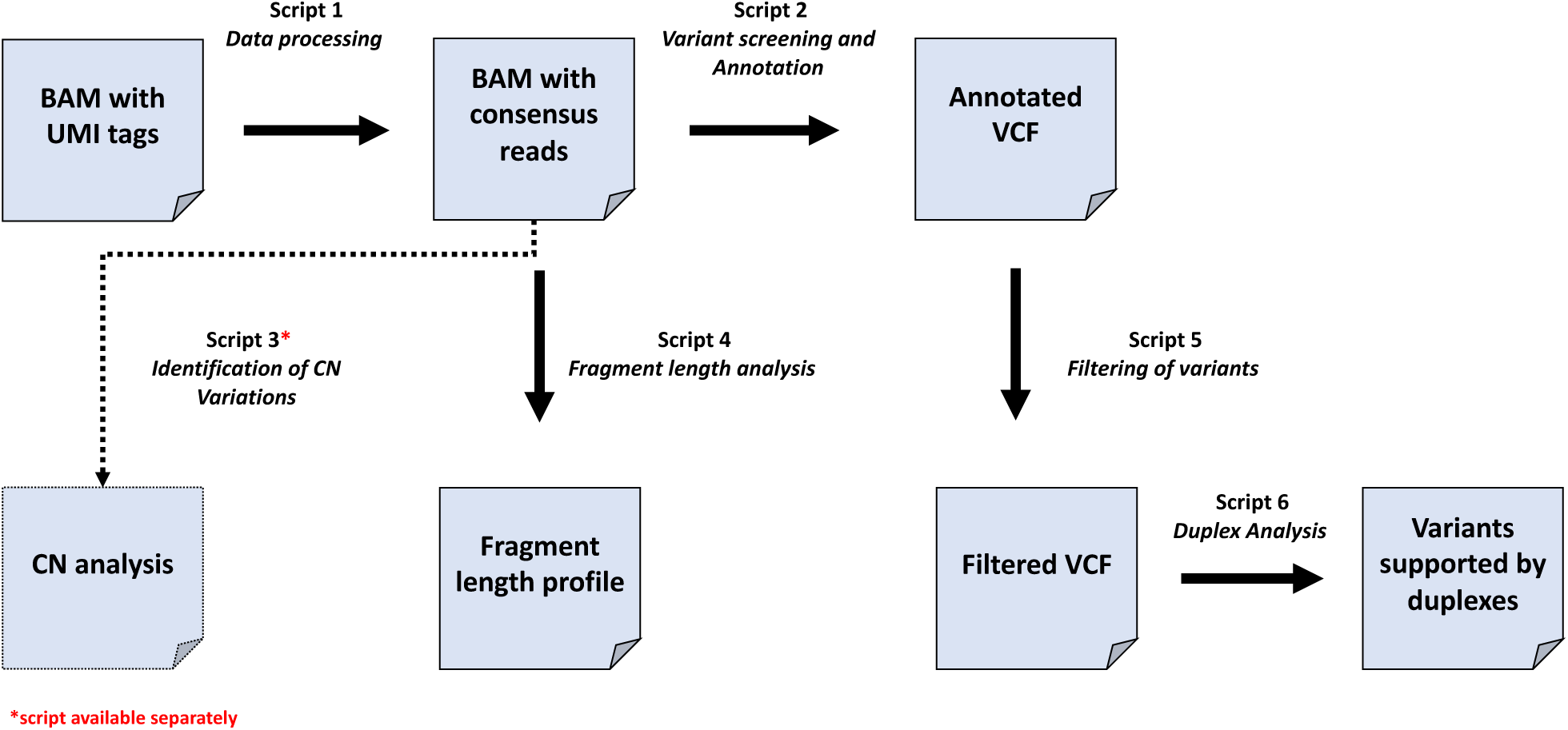
Flowchart of the developed bioinformatics pipeline.

**Supplementary Figure 2.**
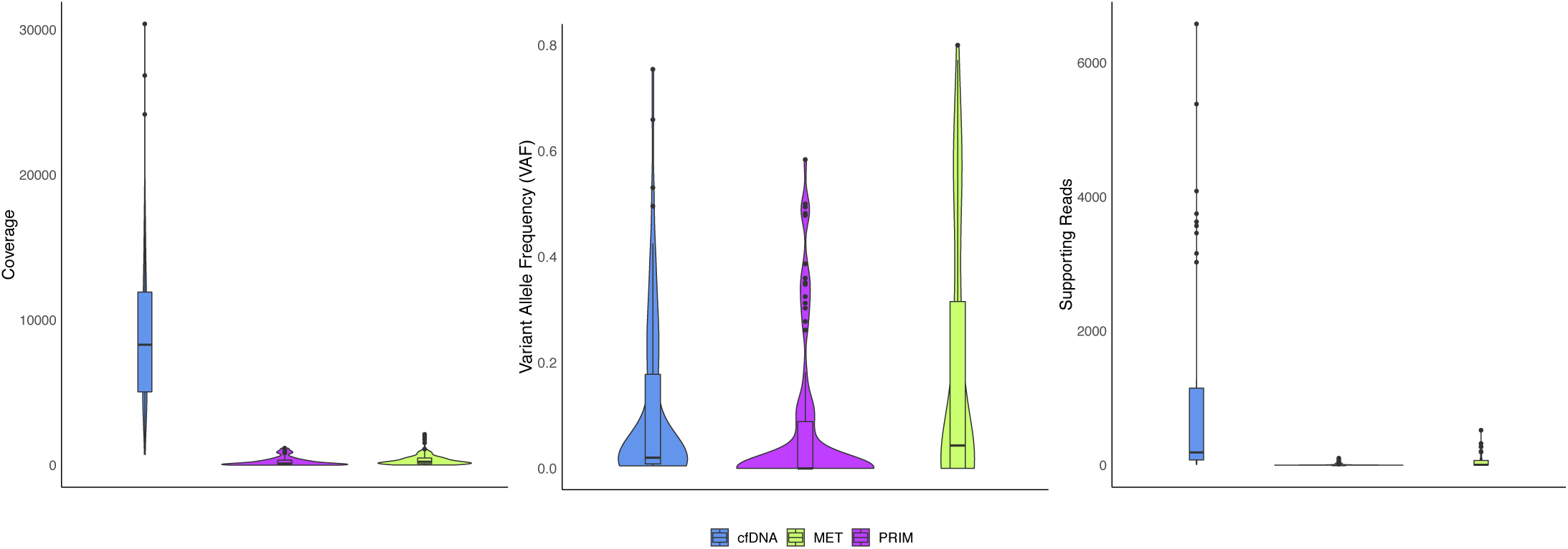
Coverage levels, VAF levels and number of supporting reads of all SNVs detected in cfDNA, primary sites and metastatic sites.

